# Age-specific dynamics of neutralizing antibodies, cytokines, and chemokines in response to La Crosse virus infection in mice

**DOI:** 10.1101/2024.05.07.592956

**Authors:** Reem Alatrash, Varun Vaidya, Bobby Brooke Herrera

**Affiliations:** Rutgers Global Health Institute, Rutgers University, New Brunswick, NJ, USA; Department of Medicine, Division of Allergy, Immunology, and Infectious Diseases and Child Health Institute of New Jersey, Rutgers Robert Wood Johnson Medical School, New Brunswick, NJ, USA

## Abstract

La Crosse virus (LACV) is a primary cause of pediatric arboviral encephalitis in the United States, particularly affecting children aged 16 years or younger. This age-related susceptibility extends to murine models, where weanling mice (3 weeks old) succumb to LACV infection, while adults (≥6 weeks old) demonstrate resistance. Despite its clinical relevance, the host immune response to LACV is not fully understood. In this study, we investigated the roles of neutralizing antibodies (nAbs), cytokines, and chemokines in weanling and adult mice following infection with 5x10^5^ plaque forming units (PFU) of LACV. We observed significant age-related differences in viral titers and survival. Weanling mice demonstrated early disease onset with elevated peripheral viremia, but passive transfer of adult serum, confirmed to have nAbs, to naïve weanlings prior to infection completely rescued them from death. Cytokine and chemokine profiling revealed distinct kinetics and age-specific immune responses. Adult mice had increased Th1 cytokines, Th9/Th17/Th22/Treg cytokines, and many chemokines. In contrast, weanlings had higher Th2 cytokines, correlating with symptoms onset. Flow cytometry and intracellular cytokine staining further demonstrated that weanling mice produced higher levels of IL-4 by CD4^+^ and CD8^+^ T cells compared to adults, regardless of infection status. Conversely, LACV-infected adult mice had increased IFN-γ production by CD8^+^ T cells compared to uninfected adults. Finally, adoptive transfer of splenocytes from immune adult mice to naïve weanlings delayed neurological symptoms and improved survival, highlighting the protective role of immune adult cells against LACV. In conclusion, this study links nAbs and cytokine and chemokine responses to protective immunity in adult mice, contrasting with the pathogenesis seen in weanlings. These findings underscore the importance of further research into innate and adaptive immune mechanisms in LACV infection.

## 1. Introduction

La Crosse Virus (LACV) is a single-stranded, negative-sense RNA virus belonging to the *Peribunyaviridae* family in the order *Bunyavirales* (1). Its transmission to humans primarily occurs through the bite of infected Eastern Tree Hole mosquitoes (*Aedes triseriatus*), although other mosquito species can harbor LACV, potentially contributing to the disease emergence (2). Upon entry into the host, LACV initially replicates in skin muscle cells, facilitating its spread into various tissues and organs (3, 4). While the majority of LACV-infected individuals manifest only mild, febrile symptoms, a subset experiences severe neurological complications, including altered behavior, seizures, coma, or fatality, as the virus invades the central nervous system (CNS) (5). Notably, severe disease and neurological manifestations of LACV infection are predominantly observed in children under the age of 16 years, indicating a higher susceptibility among this age group (6).

Vahey et al., recently reported 35-130 LACV disease cases per year using data collected in the United States between 2003 and 2019 (7). The clinical significance of LACV, coupled with its ability to also induce disease in murine models, has designated it as a prototype for studying viruses within the *Peribunyaviridae* family (8). Weanling mice (3 weeks old) exhibit susceptibility to LACV infection, whereas adult mice (≥6 weeks old) display resistance, mirroring the age-related susceptibility observed in humans (3, 8, 9). Unraveling the intricacies of host responses to LACV infection is important, particularly in delineating factors contributing to disease resistance or susceptibility. Peripheral immune control of LACV access to the CNS is believed to be crucial for disease resistance in adult mice. (10). Notably, adult mice exhibit resistance to peripheral infection, even when the virus is introduced via intraperitoneal (IP) and intradermal (ID) routes. In contrast, they are vulnerable to intracranial (IC) infection, further emphasizing the role of peripheral immunity in restricting CNS access by LACV(4, 11).

Prior investigations have shed light on the significance of both innate and adaptive immune responses following LACV infection (10, 11). Type I interferon responses have garnered considerable attention, demonstrating their indispensable role in curbing LACV replication and mitigating disease severity (12). Early cellular innate immune responses, particularly those imparted by myeloid dendritic cells, proved critical for protection in wild type adult mice infected with LACV (11). The same study also showed that treatment of young wild type mice with type I interferon, particularly IFN-β, resulted in protection mediated by myeloid dendritic cells (11). Moreover, depletion of adaptive immune cells, including both B and T cells, resulted in neurological disease in adult mice infected with LACV, suggesting that these cell types are essential for protection against neuropathology and death (10). However, the broader immune response including cytokines and chemokines remains underexplored. These signaling molecules, produced by various cell types in response to viral infections, orchestrate immune responses and exert influences on disease outcomes (13–16). Thus, deciphering the dynamics of cytokines and chemokines during LACV infection is important for unraveling their contributions to age-specific protection or pathogenesis.

Cytokines and chemokines undergo substantial modulation upon LACV infection across diverse cell types. Notably, human neuron-astrocyte co-cultures respond to LACV infection in vitro within 48 hours post infection by eliciting robust proinflammatory cytokines and chemokines, including IFN-γ, IL-6, TNF-α, IL-8, CCL4, CCL5, CXCL10 (17). In a separate study, glial cells derived from the brains of young LACV-infected mice produced significant levels of IFN-β (18). Furthermore, preliminary observations have hinted at a proinflammatory cytokine response in both the periphery and the brain following LACV infection, with elevated levels of IL-6 and IL-12p40 in infected mice, which were reported to be in comparable levels in young and adult mice (11, 19). Interestingly, administration of IL-12 or GM-CSF encoded by a plasmid prior to LACV infection in *Ifnar1^−/−^* mice was shown to increase survival rates, suggesting a potential protective role of these cytokines (20). Similarly, proinflammatory cytokines have also been reported during an Oropouche virus infection, a distantly related bunyavirus (21). Cytokines and chemokines therefore seem to act as important contributing factors in the immune response to LACV and other bunyavirus infections.

Given the contributions of cytokines and chemokines during LACV infection, this study aims to address a knowledge gap by investigating the involvement of these molecules in age-related susceptibility to LACV infection and their correlation with other components of peripheral adaptive immunity, including neutralizing antibodies (nAbs) and T cells. To address this knowledge gap, we present an in-depth analysis of nAbs, cytokine, and chemokine levels of adult and weanling mice throughout the course of infection with 5x10^5^ plaque forming units (PFU) of LACV.

## 2. Results

### 2.1 Age-related susceptibility is maintained despite infection with 5x10^5^ PFU of LACV

An unpublished observation reported no significant difference in cytokine levels between adult and weanling mice infected with 1000 PFU of LACV (11). To investigate the potential impact of infectious dose on immune system responsiveness to LACV infection, we hypothesized that a higher dose might better reveal differences in cytokine and chemokine levels if they exist. Previous studies have shown that infection with 10^5^ PFU of LACV maintains age-related susceptibility in mice (11). Peripheral inoculation routes, such as IP and ID, yield consistent age-related susceptibility and survival outcomes, with similar LACV dissemination to muscle cells near the injection site (4, 11). Based on these findings, we used IP injection for LACV infection studies. We aimed to assess the effects of an infectious dose five-fold higher than the previously reported dose by monitoring the development of neurological signs in infected mice and measuring viral titers in peripheral blood. The five-fold increase in infectious dose was chosen to reflect the higher end of the natural infectious dose transmitted by mosquito bites, while maintaining age-specific susceptibility (22). Following infection, serum from adult (n=16) and weanling (n=14) mice were serially collected for downstream processing (Fig. 1).

**Figure 1.**
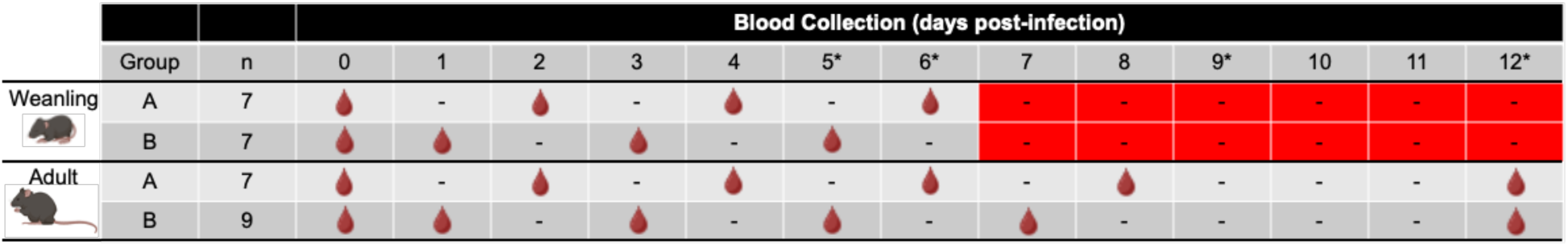
Sample collection schema. Adult and weanling mice were divided into two groups: Group A was bled on even days and Group B on odd days. Serum samples from each mouse were aliquoted for subsequent analysis of viremia, antibody neutralization, and cytokine and chemokine levels. Samples collected on day 0 served as controls. *, days when mice were removed from the study upon reaching the euthanasia endpoint. Red shaded squares, all weanling mice succumbed to infection. -, indicates no blood collection.

Injection of 5x10^5^ PFU of LACV demonstrated a similar difference in survivability between adult and weanling mice, reinforcing the age-related susceptibility to LACV infection. Adult mice exhibited markedly higher survival rates compared to weanling mice (p<0.001), with approximately 50% of adults surviving up to 12 days post-infection (dpi) (Fig. 2A). Weanling mice began exhibiting signs of neurological disease starting at 5 dpi. Our findings show that higher doses of LACV infection results in disease onset similar to that of lower infectious doses that have historically been used (Fig. 2B). Consistent with published reports, we detected LACV viremia between 1-3 dpi in both adult and weanling mice, with levels significantly higher in weanling mice at 2 dpi (Fig. 2C) (3). This observation underscores the retention of age-related susceptibility to LACV even at higher infectious doses, reaffirming the correlation between viremia and survivability.

**Figure 2.**
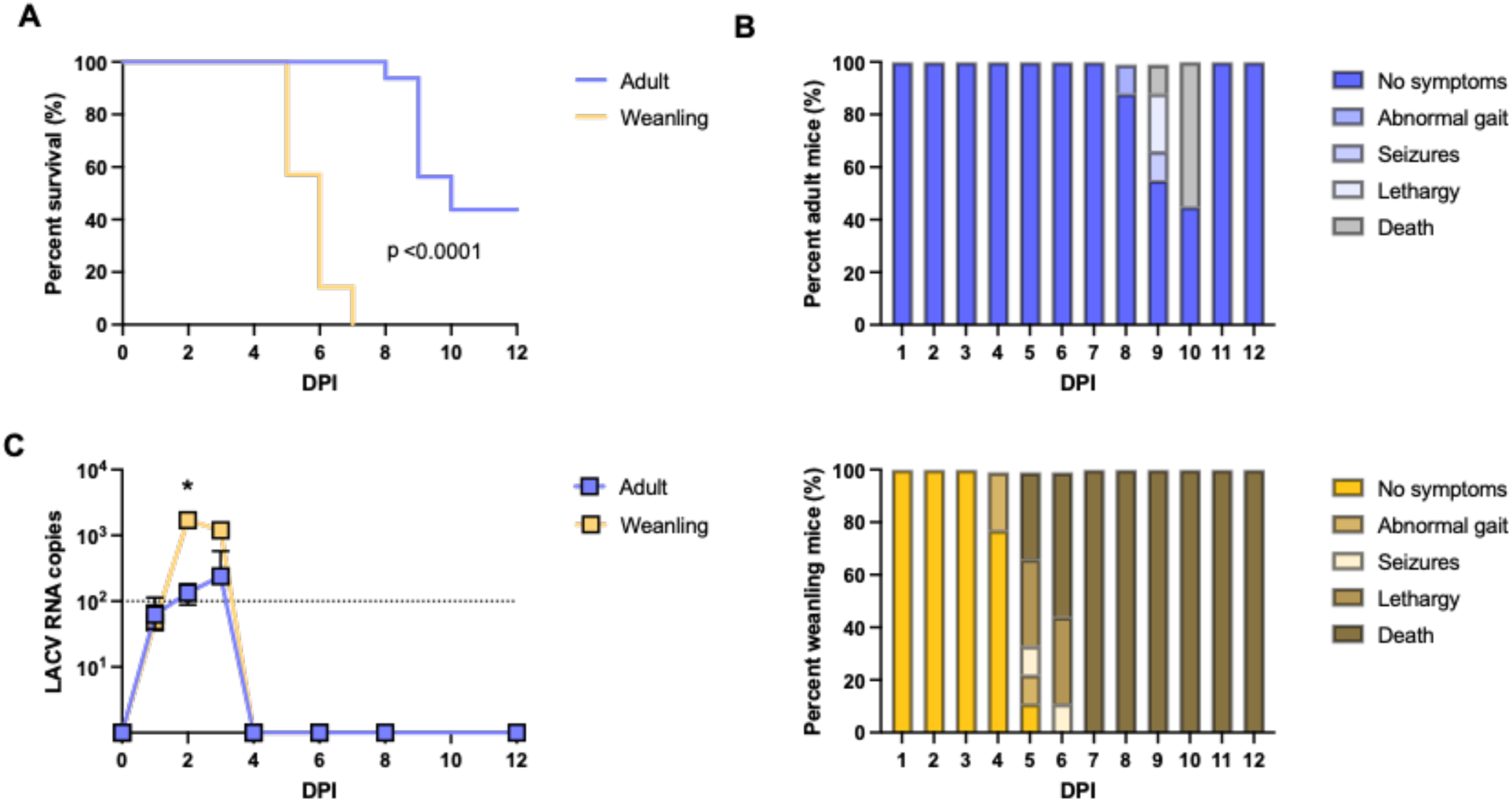
Impact of 5x10^5^ PFU of LACV infection on viremia and symptoms. **(A)** Percent survival of adult and weanling LACV-infected mice. Survival analysis was performed using the log-rank test (p<0.001) (n=7 weanling mice, n=9 adult mice). **(B)** Percent mice showing LACV-induced outcomes in adult (top) and weanling (bottom) mice. **(C)** Viral titers in sera of LACV-infected adult and weanling mice, represented by LACV RNA copies quantified by qPCR. Statistical analysis was performed using Mann-Whitney U test. **\***p<0.05 indicates significant differences. DPI, days post-infection.

### 2.2 Neutralizing antibody titers differ between LACV-infected adult and weanling mice, with the ability of adult immune serum to protect weanlings from death

Neutralizing antibody (nAb) responses are pivotal for viral clearance in many infections, yet our findings reveal an intriguing disparity between nAb titers and viral clearance in weanling mice. Following infection with 5x10^5^ PFU of LACV, nAbs peaked in adult mice between 8 and 12 dpi, whereas they peaked earlier in weanling mice between 4 and 6 dpi, coinciding with the onset of neurological signs (Fig. 3A-C). Despite the early nAb responses in weanling mice, 100% succumbed to the infection by 7 dpi (Fig. 2A). These results suggest that viral clearance and the onset of LACV-induced neurological signs may not directly correlate with the nAb responses. To assess the protective efficacy of nAbs derived from adult mice, we conducted an adoptive transfer of serum containing nAbs from previously infected adult mice into naïve weanlings. The serum transfer was performed at three distinct time points relative to LACV infection: one day prior to infection, one day post-infection, and at symptom onset (4 dpi). We observed that 100% of weanling mice that received serum one day before LACV infection survived the viral challenge (Fig. 3D). When serum was administered one day post-infection, 50% of the weanling mice survived. Notably, all weanling mice that received the serum at symptom onset succumbed to the infection, similar to the control group that received saline. These findings suggest that the early transfer of serum containing nAbs from previously infected adult mice confers a significant protective effect against LACV in weanling mice.

**Figure 3.**
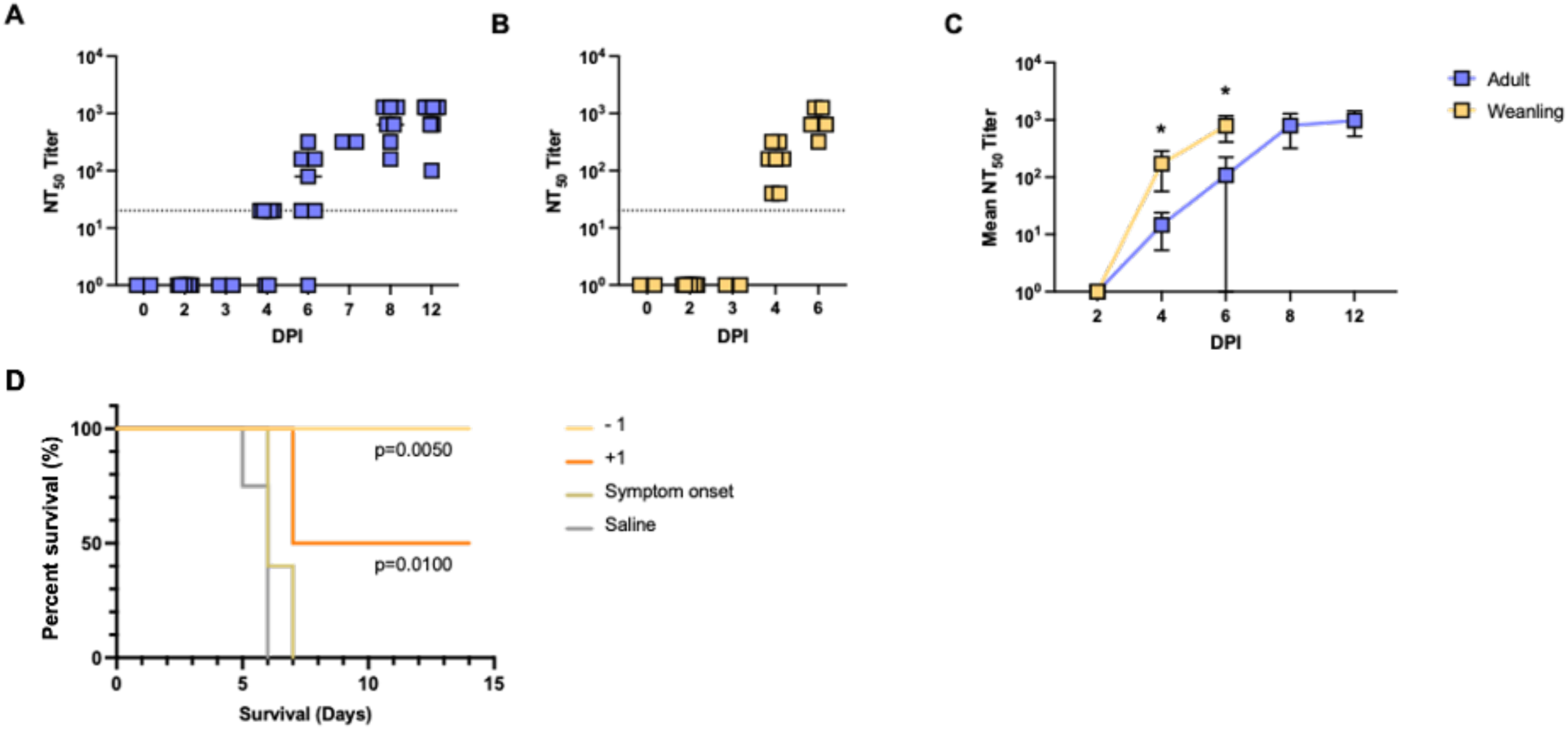
Neutralizing antibody titers differ between LACV-infected adult and weanling mice, with the ability of immune adult serum to protect weanlings from death. **(A,B)** NT_50_ titers from LACV-infected adult and weanling mice throughout the course of infection. **(C)** Mean NT_50_ titers between adult and weanling mice. Statistical analysis was performed using the Mann-Whitney U test; **\***p<0.05. **(D)** Percent survival of weanling mice following adoptive transfer of immune adult serum or saline prior or post-LACV challenge. -1, immune adult serum transferred to naïve weanling mice one day prior to LACV challenge (n=6; p=0.005). +1, immune adult serum transferred to naïve weanling mice one day post-LACV challenge (n=4; p=0.01). Symptoms onset, immune adult serum transferred to naïve weanling mice at 4 days post-infection (n=4; not significant). Saline, saline transferred to naïve weanling mice one day prior to LACV challenge as control (n=4; not significant). Survival analysis was performed using the log-rank test.

### 2.3 Cytokines and chemokines are differentially regulated in adult and weanling mice

To elucidate how serum cytokine and chemokine levels are modulated during LACV infection in adult and weanling mice, we conducted a Luminex panel capable of simultaneously detecting and quantifying Th1/Th2/Th9/Th17/Th22/Treg cytokines and chemokines. Aliquots from the same serum samples used to assess viremia and nAbs were assessed in this Luminex assay.

To delineate the potential roles of serum cytokines and chemokines in protection or pathogenesis at different infection stages, we classified the course of infection into three phases: (1) peripheral viremia (1-3 dpi), (2) symptoms onset (4-6 dpi), and (3) survival (>7 dpi). Additionally, we categorized the analyzed cytokines and chemokines into three groups: Th1/Th2 cytokines, Th9/Th17/Th22/Treg cytokines, and chemokines Among the 21 modulated cytokines and chemokines, 9 belonged to the Th1/Th2 cytokine group (GM-CSF, IFN-γ, IL-1β, IL-4, IL-6, IL-12p70, IL-13, IL-18, TNF-α, 6 were classified under the Th9/Th17/Th22/Treg cytokine group (IL-9, IL-10, IL-17A/CTLA-8, IL-22, IL-23, IL-27), and the remaining 6 were identified as chemokines in LACV-infected mice (Eotaxin/CCL11, GROα/CXCL1, IP-10/CXCL10, MCP-1/CCL2, MIP-1α/CCL3, RANTES/CCL5. Our data reveal significant fluctuations in cytokine and chemokine levels between adult and weanling mice, as shown by the heatmap illustrating p-values that compare these levels during the first 6 days of LACV infection, prior to weanling succumbing to infection (Fig. 4).

**Figure 4.**
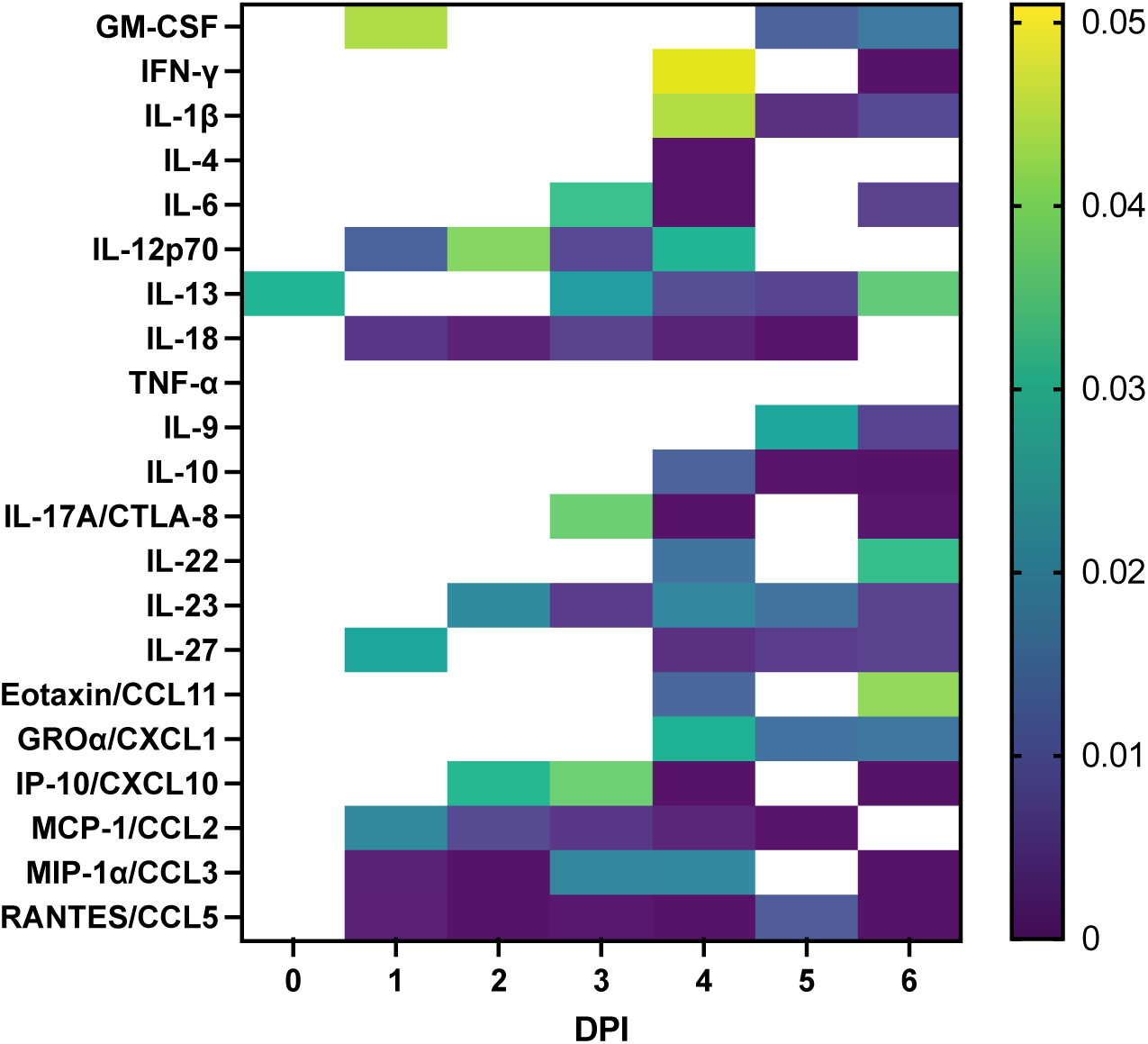
Serum cytokines and chemokines levels differ in adult and weanling mice. Heatmap of p-values derived from Mann-Whitney U test comparing cytokine and chemokine levels in adult and weanling mice during the course of LACV infection (1-6 days post-infection [DPI]). The rows represent cytokines and chemokines and the columns represent the time at which the sample was collected post-infection. Darker colors represent higher p-value.

#### 2.3.1 Dynamics of Th1/Th2 Cytokines during LACV Infection

IFN-γ, IL-18, TNF-α, and IL-13 emerged as the predominant Th1/Th2 cytokines detected in serum samples from both adult and weanling mice (Fig. 5). During the peripheral viremia phase, a significant elevation in IL-18 levels was observed as early as 1 dpi, followed by subsequent significant peaks in IL-12p70 and GM-CSF in adult mice (Fig. 5A, 5F, 5H). Notably, levels of IL-18 and IL-12p70 peaked during this early phase before gradually declining as the infection progressed. Conversely, weanling mice displayed a notable significant increase in levels of IL-6 towards the latter part of the peripheral viremia phase, one day prior to symptoms onset (Fig. 5E).

**Figure 5.**
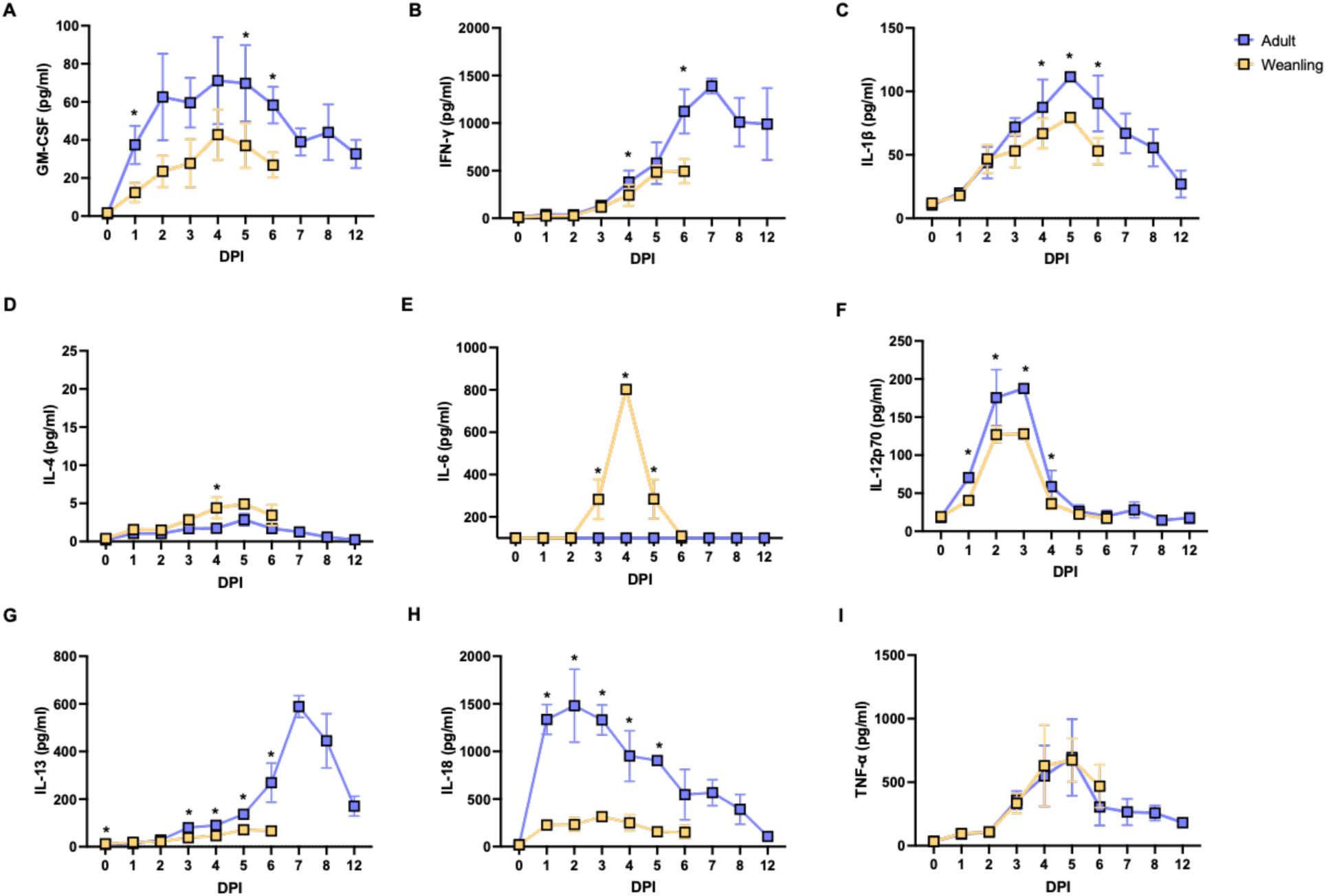
Dynamics of Th1/Th2 cytokines during LACV infection. Line charts representing levels of Th1/Th2 related cytokines (A-I) in sera of adult and weanling mice during LACV infection. Data represent average ± standard deviation. Statistical analysis was performed using the Mann-Whitney U test. **\***p<0.05 indicates significant differences.

At the symptoms onset phase, occurring between 4-6 dpi, several cytokines including IFN-γ, IL-1β, TNF-α, GM-CSF, IL-13, and IL-4 were significantly elevated in both adult and weanling mice (Fig. 5A-D, 5G, 5I). Most cytokines reached their peak levels during this phase, with adult mice exhibiting higher levels of IL-1β, TNF-α, GM-CSF, IL-13, and IFN-γ compared to weanlings. Conversely, IL-4 and IL-6 were significantly elevated in weanlings compared to adult mice (Fig. 5D-E). Finally, during the survival phase (>7 dpi) IFN-γ and IL-13 emerged as the primary cytokines specifically elevated during this late phage of infection in adult mice.

#### 2.3.2 Dynamics of Th9/Th17/Th22/Treg Cytokines during LACV Infection

Infection with 5x10^5^ PFU of LACV revealed elevated levels across all Th9/Th17/Th22/Treg cytokines analyzed in both adult and weanling (Fig. 6). Notably, IL-9, IL-10, and IL-17A emerged as the most abundant cytokines within this category throughout the infection course in both age groups. During the early peripheral viremia phase, peaks in IL-9 and IL-17A levels underscored their early involvement in the immune response to LACV infection in adult and weanling mice (Fig. 6A, 6C). Additionally, at 2 dpi, IL-23 levels were significantly higher in adult mice compared to weanlings, mirroring the kinetics of IL-17A, which exhibited significantly elevated levels in adult mice by 3 dpi (Fig. 6C, 6E).

**Figure 6.**
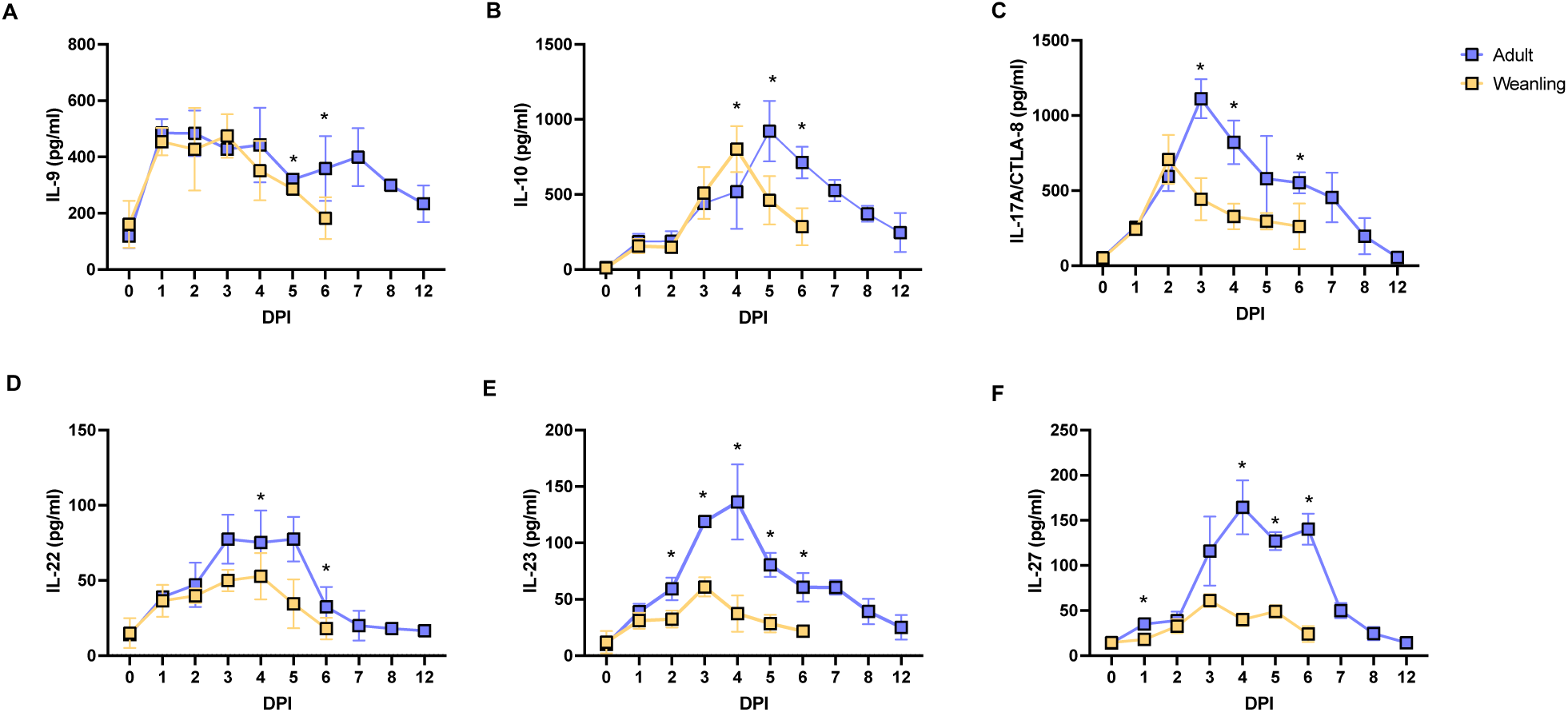
Dynamics of Th9/Th17/Th22/Treg cytokines during LACV infection. Line charts representing levels of Th9/Th17/Th22/Treg related cytokines (A-F) in sera of adult and weanling mice during LACV infection. Data represent average ± standard deviation. Statistical analysis was performed using the Mann-Whitney U test. **\***p<0.05 indicates significant differences.

During the symptoms onset phase, peaks in IL-22, IL-23, and IL-27 levels were observed, with a notable increase observed in adult mice (Fig. 6D-F). In contrast, IL-10 levels exhibited a significant increase in weanling mice at 4 dpi, followed by a rapid decline one day later, reaching levels significantly lower than those in adult mice. By 6 dpi, Th9/Th17/Th22/Treg cytokine levels gradually decreased in adult mice, ultimately returning to baseline levels.

#### 2.3.3 Dynamics of Chemokines During LACV Infection

Infection with 5x10^5^ PFU of LACV also triggered a significant increase in CXCL1 and CCL11, followed by CXCL10, in both adult and weanling mice compared to uninfected controls (Fig. 7A-C). This increase manifests as early as 1 dpi for CCL11 and CXCL1. Particularly, CCL11, CCL3, and CCL5 consistently exhibited higher levels in adult mice compared to weanlings throughout the infection course (Figure 7A, 7E, 7F). Intriguingly, CCL3 and CCL5 levels peaked during the survival phase in adult mice, suggesting sustained chemokine responses during later phases of infection.

**Figure 7.**
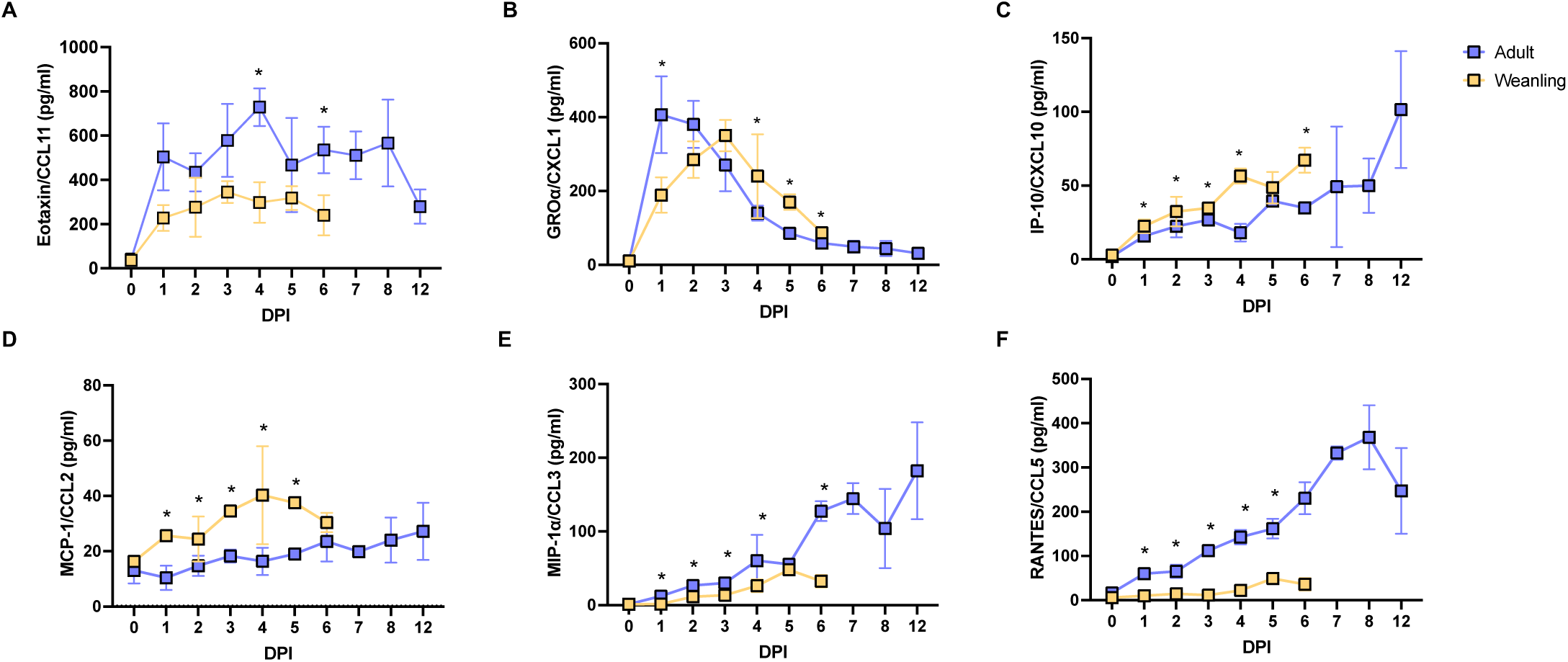
Dynamics of chemokine levels during LACV infection. Line charts representing levels of chemokines (A-F) in sera of adult and weanling mice during LACV infection. Data represent average ± standard deviation. Statistical analysis was performed using the Mann-Whitney U test. **\***p<0.05 indicates significant differences.

Conversely, distinct chemokine patterns were observed in weanling mice, with certain chemokines showing pronounced elevation whiles others exhibited minimal levels. For example, CCL2 and CXCL10 levels were consistently higher in weanling mice across all infection phases, whereas CXCL1 levels peaked exclusively during the symptom onset phase (Fig. 7B, 7D). CCL5 and CCL3 levels in weanlings remained comparable to baseline levels observed prior to infection (Figure 7E-F).

#### 2.3.4 Th1/Th2 cytokines production by CD4^+^ and CD8^+^ T cells during LACV infection

Previous studies have demonstrated the critical role of T cells in protecting adult mice from LACV-induced neurological disease, with *Rag1*^-/-^ adult mice succumbing by 15 dpi (10). Given that CD4^+^ and CD8^+^ T cells are key mediators of cytokine production in adaptive immune responses, we aimed to determine whether LACV infection induces specific cytokine expression in these T cell subsets.

We analyzed splenic CD4^+^ and CD8^+^ T cells from both adult and weanling mice at 5 dpi by flow cytometry and intracellular cytokine staining. Our results showed that weanling mice produced higher levels of IL-4 in both CD4^+^ and CD8^+^ T cells compared to adults, regardless of LACV infection status (Fig. 8A, 8D). Notably, LACV-infected weanling mice exhibited significantly higher IL-4 production in CD8+ T cells compared to uninfected controls (Fig. 8D). However, IFN-γ production showed a different pattern. While CD4^+^ T cells produced comparable levels of IFN-γ across age groups and infection statuses, CD8^+^ T cells from infected adult mice displayed increased IFN-γ production compared to uninfected adults (Fig. 8B, 8E).

**Figure 8.**
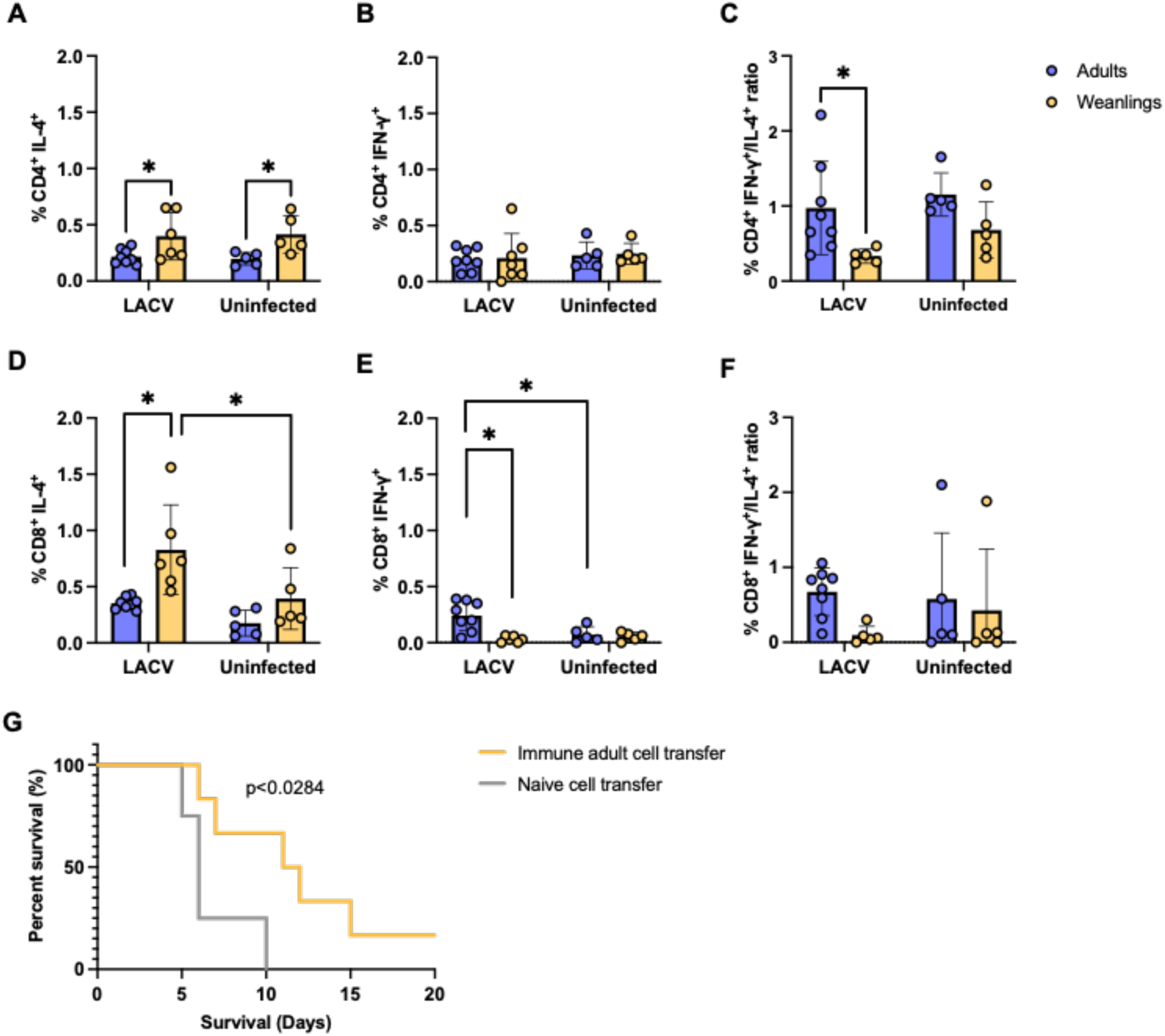
Th1/Th2 cytokine production by CD4^+^ and CD8^+^ T cells during LACV infection. **(A-F)** Bar charts showing CD4^+^ (A-B) and CD8^+^ (D-E) T cells positive for IL-4 or IFN-γ, measured by intracellular staining (ICS). IFN-γ^+^/IL-4^+^ ratios for CD4^+^ and CD8^+^ T cells are shown in (C) and (F), respectively (n=5-8). Data represent the mean ± standard deviation. Statistical analysis was performed using the Mann-Whitney U test. *p<0.05 indicates significant differences. **(G)** Percent survival of weanling mice following adoptive transfer of total splenocytes from previously infected adult mice (immune adult cell transfer), then challenged with LACV (n=6) compared to weanlings receiving cells from naïve adult mice (n=6). Survival analysis was performed using the log-rank test (p<0.0284).

To further explore the impact of LACV infection on cytokine balance, we calculated the IFN-γ/IL-4 ratio in both CD4^+^ and CD8^+^ T cells. In uninfected controls, these ratios were similar across age groups. However, in LACV-infected weanling mice, the IFN-γ/IL-4 ratio was significantly lower in CD4^+^ T cells, indicating a shift toward IL-4 production (Fig. 8C). A similar trend was observed in CD8^+^ T cells, although the difference was not statistically significant (Fig. 8F).

Given these differences in cytokine profiles between weanling and adult mice, we hypothesized that the more balanced cytokine profiles in adult mice may contribute to their greater resistance to LACV-induced disease. To test this, we adoptively transferred bulk splenocytes from immune adult mice into naïve weanlings prior to LACV infection. This transfer conferred partial protection, evidenced by improved survival rates and delayed onset of neurological disease compared to weanling controls that received naïve adult splenocytes (Fig. 8G). These findings suggest that adult T cells and the cytokine environment they establish may play a protective role in controlling LACV infection in weanling mice.

## 3. Discussion

This study presents an analysis of nAb titers and cytokine and chemokine dynamics during LACV infection, shedding light on their roles in protection and pathogenesis. Our findings demonstrate distinct patterns of cytokines and chemokines specific to age and infection phase. We observed significant increases in Th1/Th2/Th9/Th17/Th22/Treg cytokines and chemokines in response to LACV infection across both adult and weanling mice. Interestingly, we observed age-specific differences in many cytokines and chemokines throughout the course of infection (Fig. 9).

**Figure 9.**
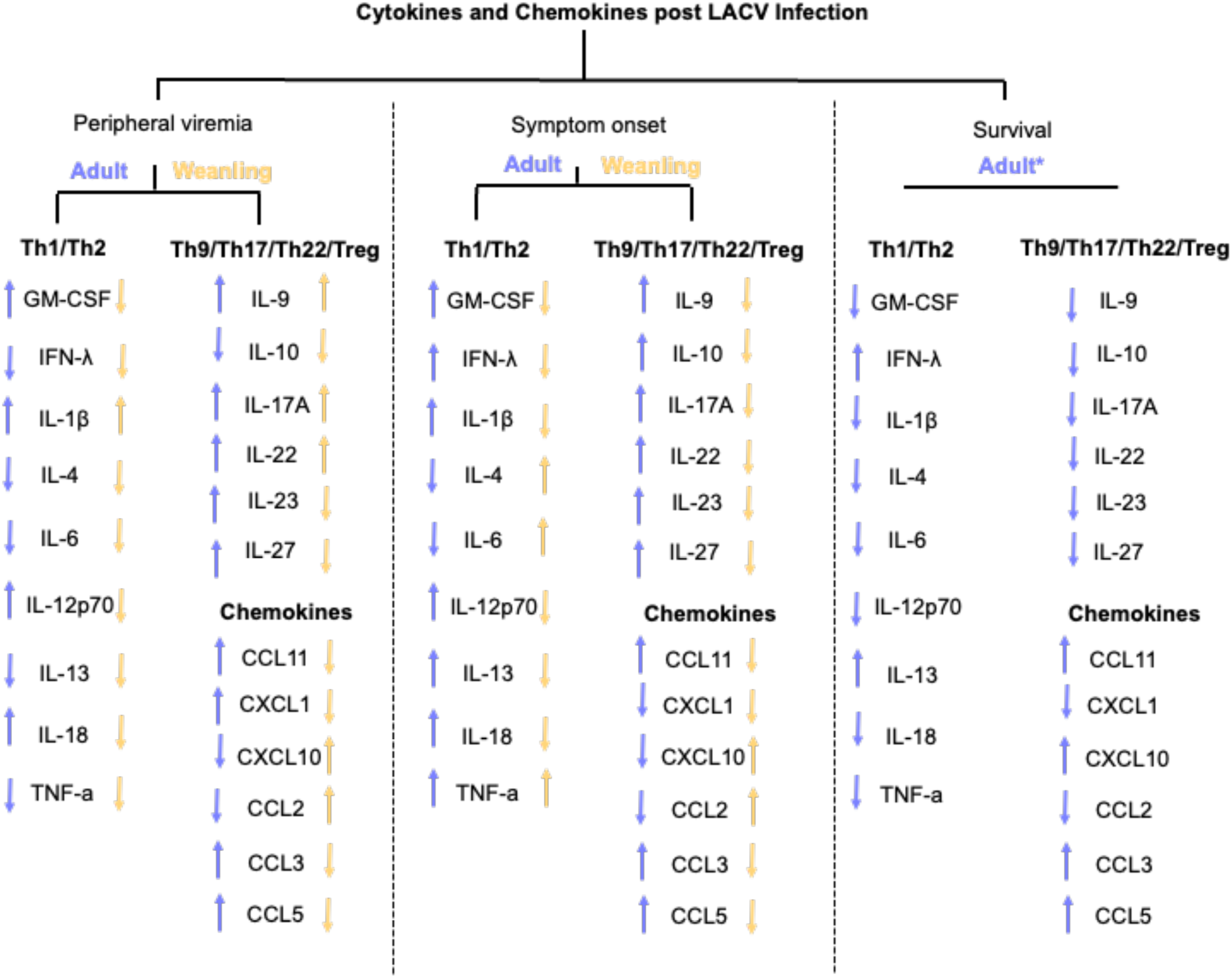
Summary of age-specific dynamic changes of cytokines and chemokines in LACV-infected adult and weanling mice. Relative shift in cytokine or chemokine levels across different stages of LACV infection, categorized as peripheral viremia, symptom onset, and survival stage, in adult and weanling mice. Purple arrows represent trends observed in adult mice, while yellow arrows represent trends observed in weanling mice. *, level compared to symptoms onset stage in adult mice.

Our data emphasize the consistent influence of age-related susceptibility, regardless of the infecting dose. Despite prior indications of age-related vulnerability using lower infectious doses of LACV, our findings expand on this concept by demonstrating that age remains a pivotal determinant of susceptibility, even when animals are infected with 5x10^5^ PFU of LACV (3). Specifically, we observed that weanling mice infected with 5x10^5^ PFU of LACV exhibited elevated peripheral viremia along with nAbs with an early onset of neurological disease. This underscores the correlation between viremia and survivability, highlighting the relevance of our model for investigating both immune response dynamics and disease pathology.

Our investigation into viral neutralization yielded unexpected findings. Despite the early production of nAbs in weanling mice, these titers failed to correlate with viral clearance, prompting speculation regarding the role of nAbs in early disease progression in weanling mice. A plausible explanation is that the virus infects the brain in weanlings by 3 dpi, rendering peripheral antibodies ineffective, which could explain the high titers observed. However, the detection of nAbs in the CNS of weanling mice by 6 dpi suggests that these antibodies can infiltrate to the CNS, where the virus is actively replicating (10). Nevertheless, these findings indicate that nAbs are ineffective in controlling the infection within the CNS as well. Notably, different neutralization capabilities have been associated with isotype switching, a phenomenon documented in other viral infections such as HIV and SARS-CoV-2 (23, 24). Interestingly, adult mice exhibited significantly lower nAb titers during the symptoms onset phase, suggesting either rapid nAbs consumption in vivo or the involvement of non-neutralizing antibodies in protection. However, given that weanling nAbs were incapable of effectively eliminating the virus in vivo, the former explanation seems less likely.

The complete protection observed in weanling mice following the early adoptive transfer of immune adult serum containing nAbs to weanlings emphasizes the importance of the humoral response in mediating protection. This finding raises several important questions, particularly regarding how adults achieve this rapid level of nAbs. One key area that remains unclear is whether the difference in protective potential between weanlings and adults is due to functional differences in the antibodies produced, the timing of their release, or a combination of both factors. Understanding these nuances is crucial, as it could shed light on the mechanisms that confer immunity and guide the development of targeted antibody-based therapies. To clarify these mechanisms, further studies focusing on antibody isotypes, function, and dynamics are necessary. While antibodies are likely key players, the protection might also be mediated by other components in the serum, such as complement or cytokines. The exact contribution of these factors remains to be clarified.

Our data demonstrate that cytokines and chemokines play a key role in age-specific susceptibility to LACV infection. We observed significant shifts in these immune mediators between adult and weanling mice during LACV infection. Given that cytokine- and chemokine-driven responses are essential for both innate and adaptive immunity, understanding their dynamics is especially important in neurotropic viral infections like LACV, which can infiltrate the CNS (25). Variability in cytokine and chemokine responses has been shown to significantly influence viral infection outcomes and the risk of severe disease(25). Additionally, previous studies have reported differences in cytokine and chemokine levels between adults and young individuals, even under healthy conditions (26). These findings suggest that age-related variations in these immune mediators could contribute to differing susceptibilities to viral infections, such as LACV. Therefore, further investigation into cytokine and chemokine profiles is warranted to better understand their role in the increased vulnerability of younger hosts to LACV infection.

In the case of LACV infection, whether increased cytokines and chemokines are a correlates of protection or pathogenesis remains to be fully investigated. However, in most cases, an increase in specific cytokines at phases of infection in susceptible hosts, such as weanling mice, may suggest a potential contribution of these pathways to susceptibility and disease pathogenesis. Conversely, the increase in cytokines and chemokines in disease-resistant hosts, such as adult mice, may indicate a protective role facilitated by these molecules.

Our study reveals distinct patterns of Th1/Th2 cytokines around symptom onset during LACV infection in weanling mice. Particularly intriguing is the role of IL-6, a key factor in determining the balance between Th1 and Th2 responses. Our data suggest that IL-6, along with IL-4, promotes Th2-related responses during symptom onset, correlating with higher nAb levels in weanling mice, supporting the Th2 skew hypothesis (27). Additionally, the substantial and sustained increase in Th1-related cytokines in adult mice underscores their protective role against disease, consistent with previous observations (11, 20).

Moreover, our findings in the context of Th1/Th2 cytokines are consistent with previous studies on CNS viral infections. For instance, elevated IL-6 levels in the CNS in infections with dengue or Japanese encephalitis viruses have been associated with various neurological manifestations and poor outcomes (28, 29). In contrast, our observation of the substantial and sustained upregulation of Th1-related cytokines, including IL-18, IFN-γ, IL-12, and GM-CSF, throughout LACV infection supports previously reported data of protective roles of GM-CSF and IL-12 in *Ifnar1^-/-^* mice and underscores their role in eliciting protective responses in mice resistant to the disease (20).

Among these cytokines, the pronounced elevation of IL-18 in adult mice stands out as the most significant shift induced by LACV. IL-18 has been demonstrated to stimulate the proliferation and differentiation of T cells toward a Th1 response and acts as a crucial cofactor in IL-12-induced IFN-γ production by natural killer (NK) and T cells (30). Moreover, IL-18 exerts its proinflammatory effects by upregulating cytokines such as IL-13, IL-1β, and TNF-α, consistent with our findings (31). Furthermore, IFN-γ emerged as one of the most robust responses, particularly increased in adult mice, especially during the survival phase. Our findings align with previous observations indicating increased IFN-γ production from neuron/astrocyte co-cultures infected with LACV in a dose-dependent manner (17). IFN-γ serves not only as a marker of Th1 CD4^+^ and CD8^+^ T cells and NK cells but also as a crucial antiviral mediator central to eliminating viruses from the CNS (32).

In our analysis of Th9/Th17/Th22/Treg cytokines, all exhibited increased levels in adult mice in response to LACV infection. Among these, IL-23 and IL-27 were among the most significantly increased. Although these cytokines have been predominantly studied in the context of autoimmune diseases, emerging evidence suggests their crucial roles during viral infections (33, 34). Specifically, IL-23 has been implicated in the control of viral infections within the brain, indicating its potential importance in host defense mechanisms against neurotropic viruses such as LACV (33). Similarly, IL-27 has been found to mediate NK cell effector functions during viral infections, highlighting its involvement in orchestrating innate immune responses against viral pathogens (34).

Chemokines, as a distinct group of immune mediators, demonstrated elevated levels in response to LACV infection, with increases observed in CCL5, followed by CCL3, in adult mice. In contrast, CCL2 peaked in weanling mice. The expression and release of CCL2 and CCL3 on cerebral endothelium suggest an essential role for these chemokines in regulating the trafficking of inflammatory cells across the blood-brain barrier (BBB) during CNS inflammation (35). Furthermore, CCL3 has been shown to contribute to the differentiation and migration of effector T cells in response to viral infection within the CNS (36). Additionally, CCL5 plays a crucial role in the homing and migration of effector and memory T cells during acute CNS infections (37). CCL2 emerges as a major player in mediating monocyte infiltration and the trafficking of inflammatory monocytes to the brain during viral encephalitis (38). Our findings, indicating increased levels of CCL2 in weanling mice, which by logic, potentially contributes to pathogenesis, potentially mediated through the recruitment of an overwhelming inflammatory response in the infected brain.

However, LACV-induced lymphocyte infiltration of CNS itself is less likely to contribute to pathogenesis according to a recent study (10). With this study being one of the few to focus specifically on LACV infections, significant gaps persist in our understanding of immune correlates and the mechanisms underlying immune-mediated viral clearance. More often than not, successful clearance of virus infection from the CNS requires the elimination of virus from the cells without causing damage to neurons. Ideally, immune-mediated virus clearance from neurons occurs mainly through early cytolytic processes, followed by non-cytolytic processes that are mediated by antibody specific for the virus and IFN-γ, which enable neurons to survive (39). Building upon this observation, a better understanding of adaptive immune responses is needed to address LACV-induced neurological disease.

Our data indicate that adaptive immune responses mediated by T cells, specifically CD4^+^ and CD8^+^ T cells, differ significantly between adult and weanling mice during LACV infection. The analysis of IL-4 and IFN-γ production by CD4^+^ and CD8^+^ T cells during LACV infection provides insights into the underlying mechanisms that contribute to differential immune responses in weanling versus adult mice. IL-4 and IFN-γ are key cytokines representing opposing arms of the immune response, with IL-4 linked to Th2 responses that promote humoral immunity and tissue repair, and IFN-γ associated with Th1 responses that drive cell-mediated immunity and viral clearance (40–43). In our study, the elevated IL-4 production in LACV infected weanling mice, particularly in CD8^+^ T cells, suggests a skewing toward a Th2-type response, which may be less effective in controlling LACV replication and mitigating disease severity. Published work have identified a subset of IL-4 producing-CD8^+^ T cells that have been described in humans in several diseases including HIV infection and *Mycobacterium tuberculosis* infection (44, 45). In contrast, the increased IFN-γ production observed in adult CD8^+^ T cells indicate a more robust Th1 response, which likely contributes to better viral control and protection from severe neurological disease. This is consistent with the observation that adult mice are generally more resistant to LACV-induced disease compared to weanling mice. Moreover, the shift in the IFN-γ/IL-4 ratio produced by CD4^+^ and CD8^+^ T cells in weanling mice toward a Th2 profile supports the notion that a balanced Th1/Th2 response is necessary for effective viral clearance and protection from severe disease.

Furthermore, the functional significance of a balanced Th1/Th2 response mediated by immune cells in LACV infection was demonstrated by the delayed disease onset and improved survival following total splenocyte transfer from previously infected adults to weanling mice. The fact that adult splenocytes were able to confer partial protection to weanling mice suggests that the enhanced IFN-γ production from adult CD8 T cells may play a critical role in mediating this protective effect. This emphasizes the need to investigate the functionality and specificity of these T cells, particularly whether they target viral antigens. However, it remains unclear whether the protective effect is primarily mediated by CD4^+^ or CD8^+^ T cells, and further detailed characterization is required to define their specific contributions.

It is important to acknowledge the limitations of our study. Characterization of cytokines and chemokines secreted into the serum may not fully capture the immune activity within the infected host, as the brain-resident immune response may differ from the systemic immune response. Additionally, we cannot definitively conclude whether a specific skew in Th1/Th2 responses occurred, as the cellular source of the cytokines detected in the serum remains unknown This study is primarily descriptive and based on observational data. While it provides valuable insights into the LACV-induced peripheral cytokine and chemokine response, experimental investigations are required to further elucidate the cell types and underlying mechanisms driving these dynamics in nAbs, cytokines, and chemokines.

In conclusion, our study contributes to a deeper understanding of immune responses to neurotropic bunyavirus infections, offering insights into potential therapeutic targets and biomarkers for disease management. Future research should focus on elucidating the underlying mechanisms driving age-related susceptibility, further characterizing the immune response dynamics, and exploring therapeutic interventions to mitigate the neurological complications associated with LACV infection. By advancing our understanding of host-virus interactions, we can develop more effective strategies for the prevention, diagnosis, and treatment of LACV-induced disease, ultimately enhancing public health preparedness against emerging viral threats.

## Methods

### Infection of mice with LACV and neurological disease progression criteria

All animal studies were approved by the Rutgers IACUC (protocol number 202300007). Wildtype (C57Bl/6) mice were purchased from Jackson Laboratories and maintained in a breeding colony at the Child Health Institute of New Jersey. LACV (La Crosse Virus, NR-540), a human isolate was purchased obtained from BEI Resources. Mice at 3 (weanling) or 6-8 (adult) weeks of age were inoculated with 5x10^5^ plaque forming units (PFU) of LACV in phosphate-buffered saline (PBS) intraperitoneally in a volume of 200 μl/mouse. Mice were observed daily for signs of neurological disease that included hunched posture, seizures, reluctance, or inability to move normally or paralysis. Animals with clear clinical signs of neurological disease were scored as clinical and euthanized immediately.

### Serum sample collection

Submandibular blood was collected into heparin coated tubes. Blood samples were centrifuged at 2000 xg for 10 minutes. After centrifugation, sera was aliquoted and stored in -80°C for subsequent analysis. Serum samples were collected from a total of 14-16 mice in each age group (Fig. 1). Those mice were divided into two subgroups, with each group being bled every other day. Accordingly, serum samples were collected on 0, 1, 2, 3, 4, 5, 6, 7, 8 and 12 dpi. This study design was implemented to reduce the number of mice used for ethical purposes and to ensure that the serum sample volume from each mouse remained within ethical guidelines. Serum samples collected on odd days were pooled by gender. The sample size decreased over time as some mice did not survive the infection.

To assess the correlation between cytokine and chemokine levels and indicators of viral clearance and neutralizing antibody responses, each serum sample was used in three assays: viremia, neutralization, and cytokine/chemokine quantification.

### Neutralizing antibody detection

For quantification of neutralizing antibodies, sera samples were serially diluted in quadruplicates and mixed with an equal volume of LACV corresponding to 100 PFU per in a final volume of 100 μl in DMEM/10% FBS/1% Pen Strep. The mixture was preincubated for 2 h at 37°C. The 100 μl mixtures were then added to confluent Vero cells in a 96-well plate and incubated for 3 days. After incubation, plasma/virus mixture was removed, and cells were stained with 0.1% crystal violet for 2 h. After staining, cells were rinsed with deionized water. Neutralizing antibody titers were determined by the highest plasma dilution protecting 50% of the infected wells.

### qPCR

Viral titers were quantified by qPCR. Total viral RNA was extracted from serum using Pure link viral RNA mini kit (Invitrogen) following manufacturer’s protocol. cDNA was prepared from RNA samples using the SuperScript IV VILO reverse transcription kit (Invitrogen) following the manufacturer’s instructions. cDNA samples were diluted fivefold in Rnase-free water and those diluted samples were analyzed by qPCR using PowerTrack SYBR Green kit (Appliedbiosystems) to detect LACV RNA. The primer sequences used in this study are: forward (ATTCTACCCGCTGACCATTG), and reverse (GTGAGAGTGCCATAGCGTTG). CT values were measured using a QuantStudio 3 Flex (Applied Biosysterms) machine. QuantStudio Real-Time PCR software was used to extract the data.

### Adoptive serum transfer

Adult mice previously infected with LACV were sacrificed at 14 days dpi, and serum was collected via cardiac puncture. For the adoptive transfer experiments, weanling mice received 200 µL of serum intraperitoneally at three distinct time points: one day before LACV challenge, one day after LACV challenge, or on the day of symptom onset (4 dpi). As a control, a separate group of weanling mice received 200 µL of sterile saline intraperitoneally at each of the corresponding time points.

### Bead-based multiplex immunoassay (Luminex)

To quantify the levels of cytokines in LACV infected serum samples, a magnetic bead-based multiplex immunoassay was performed using a MAGPIX® (Luminex Corporation, USA) instrument system. A mouse cytokine magnetic 26-plex panel (Invitrogen) was utilized to determine immune responses mediated through cytokine and chemokine such as (1) Th1/Th2: GM-CSF, IFN-γ, IL-1 beta, IL-2, IL-4, IL-5, IL-6, IL-12p70, IL-13, IL-18, TNF alpha, and (2) Th9/Th17/Th22/Treg: IL-9, IL-10, IL-17A (CTLA-8), IL-22, IL-23, IL-27, (3) Chemokines: Eotaxin (CCL11), GRO alpha (CXCL1), IP-10 (CXCL10), MCP-1 (CCL2), MCP-3 (CCL7), MIP-1 alpha (CCL3), MIP-1 beta (CCL4), MIP-2, RANTES (CCL5) following the manufacturer’s instructions.

### Tissues processing for flow cytometry and intracellular staining (ICS)

Spleens from LACV-infected and uninfected mice were harvested at 5 dpi, as described previously (10). Spleens were processed into single-cell suspensions by passing them through a 70 µM mesh filter, followed by red blood cell lysis using RBC Lysis Buffer (Thermo Fisher Scientific). Isolated cells were then seeded at a density of 2 x 10⁶ cells per well in 96-well U-bottom plates and cultured. Cells stimulated with PMA served as a positive control, and Brefeldin A was added during the 4-hour incubation. After incubation, cells were stained for viability and Fc receptors were blocked. Cells were stained for extracellular markers (CD8a, CD4, and CD3), fixed and permeabilized, and stained for intracellular cytokines (IL-4 and IFN-γ). Flow cytometry was performed using the Aurora system (Cytek® Biosciences), and data were analyzed with FlowJo software, gating on live cells and excluding doublets.

### Adoptive transfer of total splenocytes

Donor mice are euthanized at 14 dpi, spleens are harvested, and splenocytes are generated as described above. Naïve recipient mice receive transfusions of 10^6^ total cells in 100 ul PBS via tail vein injection and are subsequently challenged 1 day post transfusion with 10^3^ PFU LACV via intraperitoneal injection.

### Statistical analysis

All statistical analyses were performed using Prism software Version 7.01 (GraphPad) and are described in the figure legends.

## Data Availability

The data that support the findings of this study are available within the manuscript or upon request from the corresponding author.

## Acknowledgements

We would like to thank Rutgers Global Health Institute, Rutgers Robert Wood Johnson Medical School, and the Child Health Institute of New Jersey for their support. We would also like to thank the Foundation for Health Advancement for grant support (IFHA 17-24). The funders had no role in the design, data collection, data analysis, and reporting of this study.

